# Spatial separation provides the force-coupling mechanism for the spindle assembly checkpoint

**DOI:** 10.64898/2026.09.24.754109

**Authors:** Chyi Wei Chung, Daniel Grant, Danny B. H. Gold, Jack Douglas, Manton Leung, Francis A. Barr, Ulrike Gruneberg

## Abstract

The spindle assembly checkpoint (SAC) prevents anaphase until chromosomes have formed correct bipolar attachments to the mitotic spindle. Spatial separation of pericentromeric Aurora B from outer-kinetochore substrates explains how tension-dependent biorientation stabilises microtubule-kinetochore attachments. How this geometry controls the SAC and is ultimately sensed by the key checkpoint kinase MPS1 has remained unclear. Here, we show that MPS1 recruitment is regulated by the spatial relationship between pericentromeric Aurora B-INCENP and the outer kinetochore. Shortening INCENP reduces MPS1 recruitment, whereas extending INCENP delays MPS1 removal and SAC silencing. Partial NDC80 depletion prevents normal Aurora B-kinetochore separation while retaining MPS1-dependent checkpoint signalling. Conversely, acute recruitment of Aurora B to microtubule-attached outer kinetochores under tension bypasses spatial separation and rapidly restores MPS1 and downstream SAC signalling. We propose that error correction and MPS1-dependent SAC signalling share an upstream spatial-sensing mechanism in which force-dependent separation from Aurora B favours attachment stabilisation while limiting MPS1 recruitment.

## Introduction

Chromosome segregation in mitosis depends on the bipolar attachment of microtubules from opposite spindle poles to the outer kinetochores of duplicated sister chromatids (Musacchio and Desai, 2017). The fidelity of chromosome segregation is ensured by two coordinated processes. Spindle assembly checkpoint (SAC) signalling from kinetochores on non-bioriented chromosomes inhibits cell-cycle progression until all chromosomes are correctly attached to the spindle (McAinsh and Kops, 2023; Musacchio, 2015). Error correction destabilises kinetochore–microtubule attachments that fail to reach a bioriented and fully tensioned state, and thus regenerates checkpoint active kinetochores (DeLuca et al., 2006; Tanaka and Zhang, 2022). The conserved mitotic kinases Aurora B and MPS1, which localise to the pericentromere and kinetochore, respectively, are required for both these processes (Krenn and Musacchio, 2015; Pachis and Kops, 2018). Despite intense study of these kinases, how they cooperate to translate attachment state into coordinated control of error correction and checkpoint signalling remains unclear.

Restriction of Aurora B to the pericentromeric heterochromatin is a crucial determinant for error correction. This requires the chromosomal passenger complex (CPC) of which Aurora B is the catalytic core (Carmena et al., 2012). The CPC attaches to pericentromeric nucleosomes by a targeting module composed of borealin, survivin, and the N-terminus of INCENP, while Aurora B binds to and is activated by the C-terminal IN-box of INCENP (Gireesh et al., 2025; Jeyaprakash et al., 2007; Klein et al., 2006; Ruza et al., 2025; Sessa et al., 2005). These targeting and catalytic elements are linked by the long, flexible INCENP arm (Samejima et al., 2015). During spindle assembly this pool of Aurora B promotes error correction by phosphorylating outer-kinetochore substrates required for microtubule binding, thereby destabilising inappropriate attachments (Krenn and Musacchio, 2015; Welburn et al., 2010). NDC80, which forms part of the microtubule binding site in the outer kinetochore is a key component regulated in this way. Aurora B-dependent phosphorylation of the NDC80 N-terminal tail reduces its microtubule binding affinity, thus promoting error correction (Cheeseman et al., 2006; Ciferri et al., 2008; DeLuca et al., 2011; Guimaraes et al., 2008). Seminal work by Lampson, Lens and colleagues showed that microtubule-generated forces increase the distance between pericentromeric Aurora B and its outer-kinetochore substrates, thus reducing their phosphorylation and allowing kinetochore-microtubule attachments to stabilise (Lampson and Cheeseman, 2011; Liu et al., 2009). Conversely, artificially repositioning Aurora B towards kinetochores destabilised microtubule-kinetochore attachments (Liu et al., 2009). Together these findings provided a physical framework explaining the coupling of Aurora B activity to chromosome biorientation.

An important insight from these and other studies is that Aurora B indirectly promotes SAC activity through its role in error correction. In addition, Aurora B is also directly required for recruitment of MPS1 to kinetochores and thus acts to integrate the SAC with error correction (Hayward et al., 2019b; Hayward et al., 2022; Nijenhuis et al., 2013; Santaguida et al., 2011; Saurin et al., 2011). Interestingly, at kinetochores, MPS1 also contributes to microtubule turnover and chromosome biorientation, although how this function is integrated with Aurora B activity has not been fully established yet (Hayward et al., 2022; Hewitt et al., 2010; Jelluma et al., 2008; Maciejowski et al., 2017; Maure et al., 2007; Santaguida et al., 2010).

How microtubule-binding controls MPS1 activity and the role of Aurora B is therefore more complex than it might seem due to these direct and indirect effects, and different proposals have therefore been put forward to explain the mechanism (Hiruma et al., 2015; Ji et al., 2015; Khodjakov and Pines, 2010; Nezi and Musacchio, 2009). We have shown that MPS1 localisation and activity at kinetochores during checkpoint signalling is controlled by the balance of Aurora B and counteracting phosphatase PP2A-B56 (Hayward et al., 2022). Crucially, MPS1 and microtubules can bind to kinetochores simultaneously, albeit transiently, as the checkpoint signal is initiated during error correction (Hayward et al., 2022; Pleuger et al., 2024). These findings suggest that a microtubule-dependent process modulates Aurora B activity to control MPS1 recruitment indirectly, rather than directly through microtubule binding or competition with microtubules.

We now test if the spatial separation model, which was established for Aurora B-mediated error correction, can also explain how MPS1 is regulated during checkpoint signalling. Central to this model is the long, flexible arm of INCENP (“dog leash”), which connects the pericentromeric targeting module of the CPC to the Aurora B-binding IN-box and defines the potential range over which pericentromere-localised Aurora B can act (Krenn and Musacchio, 2015; Samejima et al., 2015; Santaguida and Musacchio, 2009). We show that shortening INCENP reduces MPS1 recruitment to kinetochores, and conversely that extending INCENP delays MPS1 removal and SAC silencing. Using the tuneable Halo-PROTAC system we find that partial NDC80 depletion prevents normal Aurora B-kinetochore separation but leaves sufficient NDC80 to sustain an MPS1-dependent checkpoint arrest. Furthermore, acute recruitment of Aurora B to attached outer kinetochores rapidly restores MPS1 and downstream SAC signalling, before detectable loss of stable attachments or inter-kinetochore stretch. Microtubule-generated forces therefore separate outer-kinetochore substrates from pericentromeric Aurora B, favouring attachment stabilisation while limiting Aurora B-dependent MPS1 recruitment. Together, these results indicate that error correction and MPS1-dependent SAC signalling share a common upstream force-sensing mechanism.

## Results and Discussion

### MPS1 kinetochore localisation correlates with Aurora B and NDC80 proximity

Aurora B kinase activity is necessary for MPS1 recruitment to the outer kinetochore (Hayward et al., 2019b; Hayward et al., 2022; Nijenhuis et al., 2013; Ruza et al., 2025; Saurin et al., 2011). Pericentromeric Aurora B and the outer kinetochore protein NDC80 overlap in prometaphase and become separated in metaphase (Ruza et al., 2025). We therefore asked whether this spatial relationship correlates with MPS1 kinetochore localisation. Using structured illumination microscopy (3D SIM) and fluorescence lifetime-based stimulated emission depletion (TauSTED) microscopy, we analysed HCT116 NDC80-HaloTag MPS1-mStayGold cells. In prometaphase, sister kinetochores defined by NDC80-HaloTag were separated by 0.59 ± 0.21 µm, while Aurora B formed a compact signal between the two NDC80 peaks. The mean Aurora B-NDC80 peak distance was 0.28 ± 0.03 µm, and MPS1 colocalised with NDC80 (Figure 1A, C, E, F). In metaphase, the NDC80-NDC80 distance increased to 1.27 ± 0.20 µm and the Aurora B-NDC80 peak distance to 0.63 ± 0.16 µm, while MPS1 was no longer detectable at kinetochores (Figure 1B, D, E, F). Thus, increased separation of Aurora B from NDC80 during prometaphase and metaphase accompanies MPS1 loss from metaphase kinetochores.

**Figure 1.**
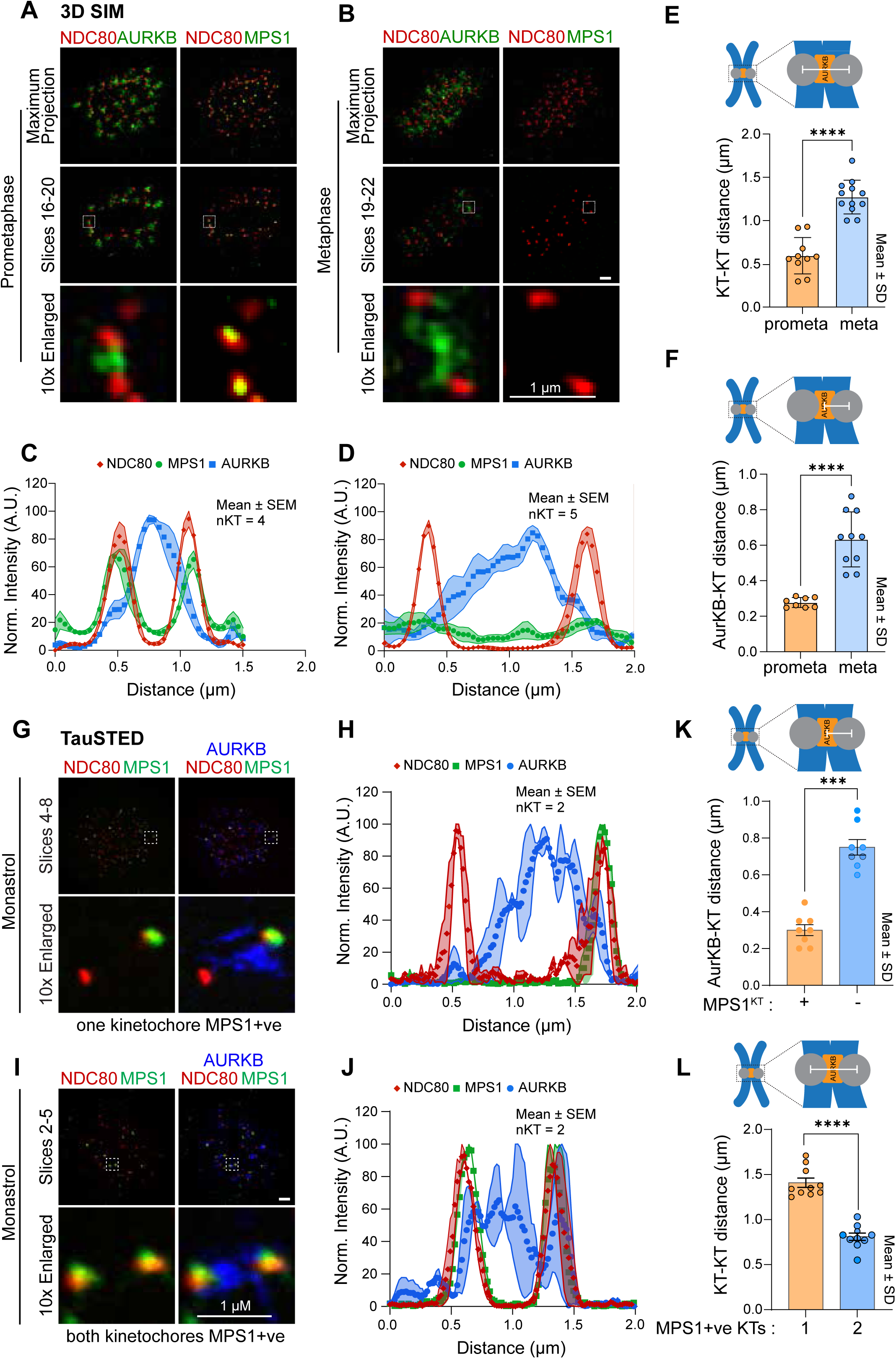
MPS1 kinetochore localisation correlates with Aurora B and NDC80 proximity. (A) HCT116 NDC80-HaloTag MPS1-StayGold cells in prometaphase or (B) metaphase were analysed by 3D SIM. NDC80-Halo was stained with JFX554 dye, and Aurora B (AURKB) was visualised by antibody staining. Representative maximum intensity projections and selected slices are shown. The indicated kinetochore pairs are shown 10x enlarged at the bottom. (C) and (D) Line scans show signal intensity across kinetochore pairs (mean ± SEM, n = 4 or 5, as indicated in the figure). (E) KT-KT distances in prometaphase and metaphase are plotted (mean ± SD; ****indicates p<0.0001). (F) Aurora B-KT distances, determined by the distance between fitted Gaussian peaks, in prometaphase and metaphase are plotted (mean ± SEM; ****indicates p<0.0001). (G) TauSTED imaging of HCT116 NDC80-HaloTag MPS1-StayGold cells arrested with monastrol overnight. Aurora B and NDC80-HaloTag were visualised as in (A). Representative maximum intensity projections of selected slices of kinetochore pairs are shown. The indicated kinetochore pair with one MPS1-positive kinetochore is shown 10x enlarged at the bottom. (H) A line scan shows signal intensity across kinetochore pairs (mean ± SEM, n = 2). (I) TauSTED imaging as in (G) but of a kinetochore pair with two MPS1-positive kinetochores. (J) Line scan as in (H). (K) Aurora B-KT distances between MPS1-positive and MPS1-negative kinetochores are plotted (mean ± SD; **** indicates p<0.0001). (L) KT-KT distances between kinetochore pairs with one or two MPS1-positive kinetochores are plotted (mean ± SEM; **** indicates p<0.0001). Paired t-tests were used for statistical analysis.

A similar relationship was evident within individual kinetochore pairs in monastrol-arrested cells. In cases where only one sister kinetochore was MPS1-positive, Aurora B was distributed asymmetrically towards that kinetochore, with a shorter Aurora B-kinetochore distance at the MPS1-positive kinetochore than at its MPS1-negative sister (0.30 ± 0.08 µm versus 0.75 ± 0.12 µm; Figure 1G, H, K). In the less frequent pairs in which both kinetochores were MPS1-positive, Aurora B distribution was more symmetric, and the kinetochore-kinetochore distance was reduced to 0.81 ± 0.13 µm (Figure 1I, J, L). Together, these observations reveal a close correlation between Aurora B proximity and MPS1 kinetochore occupancy and prompted us to test whether altering the effective reach of Aurora B changes SAC signalling.

### INCENP length defines the spatial range of Aurora B-dependent MPS1 recruitment

Stable tethering of the CPC to pericentromeric nucleosomes is essential for efficient MPS1 kinetochore recruitment (Ruza et al., 2025). In addition to concentrating Aurora B at pericentromeres this restricts its action to substrates, through the long single alpha-helix (SAH)/coiled-coil region of INCENP which acts as a spacer between the centromere-targeting module and the Aurora B-binding IN-box (Samejima et al., 2015; Vader et al., 2007). We therefore asked whether INCENP length defines the spatial range of Aurora B-dependent MPS1 recruitment. To do this we generated HeLa Flp-In TRex cells expressing endogenously tagged MPS1-GFP as well as mScarlet-INCENP^FL^, truncated mScarlet-INCENP^ΔSAH^ or mScarlet-INCENP^Δ137-789^, the latter lacking both the SAH and much of the intervening unstructured regions connecting the centromere targeting domain and the IN-box (Figure 2A). Full-length INCENP and both truncations bound borealin and survivin and associated with T-loop-phosphorylated Aurora B (pT232) at comparable levels (Figure 2B). Histone H3 Ser10 phosphorylation was also maintained (Figure S1A-D). Following depletion of endogenous INCENP, the transgenes were induced. The mSc-INCENP constructs were expressed at similar levels to one another, although less than endogenous INCENP, and importantly localised to pericentromeric chromatin (Figure 2C-E). Under these conditions, full-length mSc-INCENP supported robust MPS1 kinetochore localisation, whereas the number and intensity of MPS1-positive kinetochores was decreased in cells expressing mSc-INCENP^ΔSAH^ and further reduced with mSc-INCENP^Δ137-789^ (Figure 2D, F). Removing microtubules with nocodazole partially alleviated the MPS1 localisation defect of the shortened INCENP variants (Figure S1E-G). This is consistent with the idea that, when microtubule-dependent pulling forces are absent, the reduced distance between pericentromeric chromatin and the outer kinetochore partly compensates for the shorter INCENP length. Cells expressing mSc-INCENP^ΔSAH^ or mSc-INCENP^Δ137-789^ also failed to maintain a robust taxol-induced mitotic arrest, although the phenotype was less severe than MPS1 inhibition, indicating residual SAC activity (Figure 2G-I). In unperturbed mitosis, this residual activity was sufficient for many cells to progress through mitosis with normal or moderately prolonged timing and without overt defects in chromosome biorientation (Figure S2A-E), consistent with previous observations (Vader et al., 2007).

**Figure 2.**
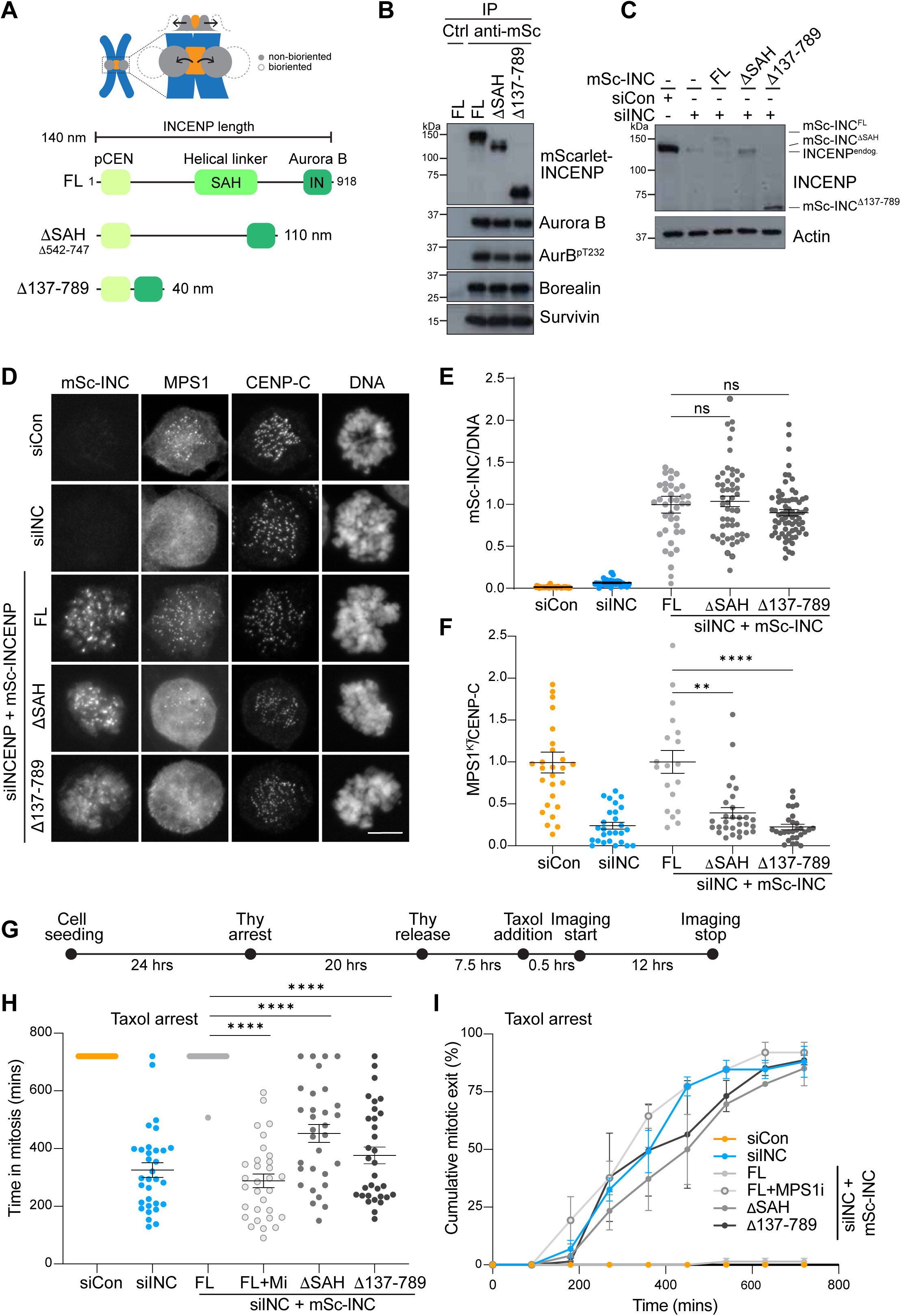
INCENP length defines the spatial range of Aurora B-dependent SAC signalling. (A) Schematic of the different mScarlet-tagged INCENP truncations. (B) Mitotically arrested HeLa Flp-In TREx cells expressing the indicated INCENP transgenes were lysed, and the different versions of INCENP were immunoprecipitated with anti-mCherry antibodies (cross-reacting with mScarlet). The immunoprecipitates were Western blotted with the indicated antibodies. (C) HeLa Flp-In TREx cells depleted of the endogenous INCENP and expressing the indicated INCENP transgenes were lysed and blotted with antibodies against INCENP, and actin as a loading control. (D) HeLa Flp-In TREx cells, depleted of endogenous INCENP and expressing endogenously GFP-tagged MPS1 as well as the indicated INCENP transgenes were fixed and stained as indicated. Prometaphase cells at equivalent stages of mitosis were analysed. Scale bar, 10 µm. (E) Quantitation of the mScarlet-INCENP signal in (D) showing mean ± SEM for siControl, n = 44; siINCENP, n = 60; siINCENP + FL, n = 39; siINCENP + ΔSAH, n = 52; siINCENP + Δ137-789, n = 70; ns, p > 0.999; Kruskal-Wallis test was used. (F) Quantification of kinetochore (KT) MPS1 normalised to kinetochore CENP-C. siControl, n = 27; siINCENP, n = 27; siINCENP + FL, n = 19; siINCENP + ΔSAH, n = 29; siINCENP + Δ137-789, n = 28). Kruskal-Wallis test was used for statistical analysis, where *** indicates p<0.001; **** p<0.0001. (G) Time line of the experiment. HeLa Flp-In TREx cells depleted of the endogenous INCENP and expressing the indicated INCENP transgenes were synchronised in early S-phase with 2 mM thymidine, released for 8 hrs and treated with 0.1 µM Taxol for 30 min before imaging. 2 µM MPS1 inhibitor AZ3146 was added to mScarlet-INCENP^FL^ cells as a control. (H) and (I) Quantification of (G). Graphs show repeat-averaged values for 3 biological repeats (n>10 per repeat), normalised to FL-INCENP. Kruskal-Wallis test was used for statistical analysis, with **** p<0.0001, *** p<0.001, ** p<0.01, * p<0.05.

If the effective reach of centromere-localised Aurora B contributes to MPS1 recruitment, extending INCENP should have the opposite effect and delay the loss of Aurora B-dependent signalling as microtubule-generated forces develop. We therefore lengthened INCENP by inserting two SAH domains, using the previously described 8KQ variant to reduce microtubule binding, referred to as INCENP^+2x8KQ^ (van der Horst et al., 2015). A single SAH is predicted to extend INCENP by approximately 80 nm, giving the INCENP^+2x8KQ^ construct a potential increase in reach of up to 160 nm (Samejima et al., 2015) (Figure 3A). Before analysing spindle checkpoint function, we first confirmed that mSc-INCENP^FL^ and mSc-INCENP^+2x8KQ^ were expressed at similar levels, localised to centromeric chromatin in early mitosis, and supported comparable prometaphase MPS1 kinetochore levels (Figure 3B-E). Control cells expressing full-length INCENP entered anaphase 57 ± 8 min after nuclear envelope breakdown (NEBD). In contrast, INCENP^+2x8KQ^ cells frequently remained with largely aligned chromosomes for extended periods and only entered anaphase after 173 ± 68 min on average (Figure 3F, G). MPS1 inhibition abolished the cell cycle delay, showing that it was SAC dependent (Figure 3G). Consistent with impaired spindle checkpoint silencing, INCENP^+2x8KQ^ cells arrested with the proteasome inhibitor MG132 showed a modest but significant increase in kinetochore MPS1 at metaphase in comparison to control cells (Figure 3H, I).

**Figure 3.**
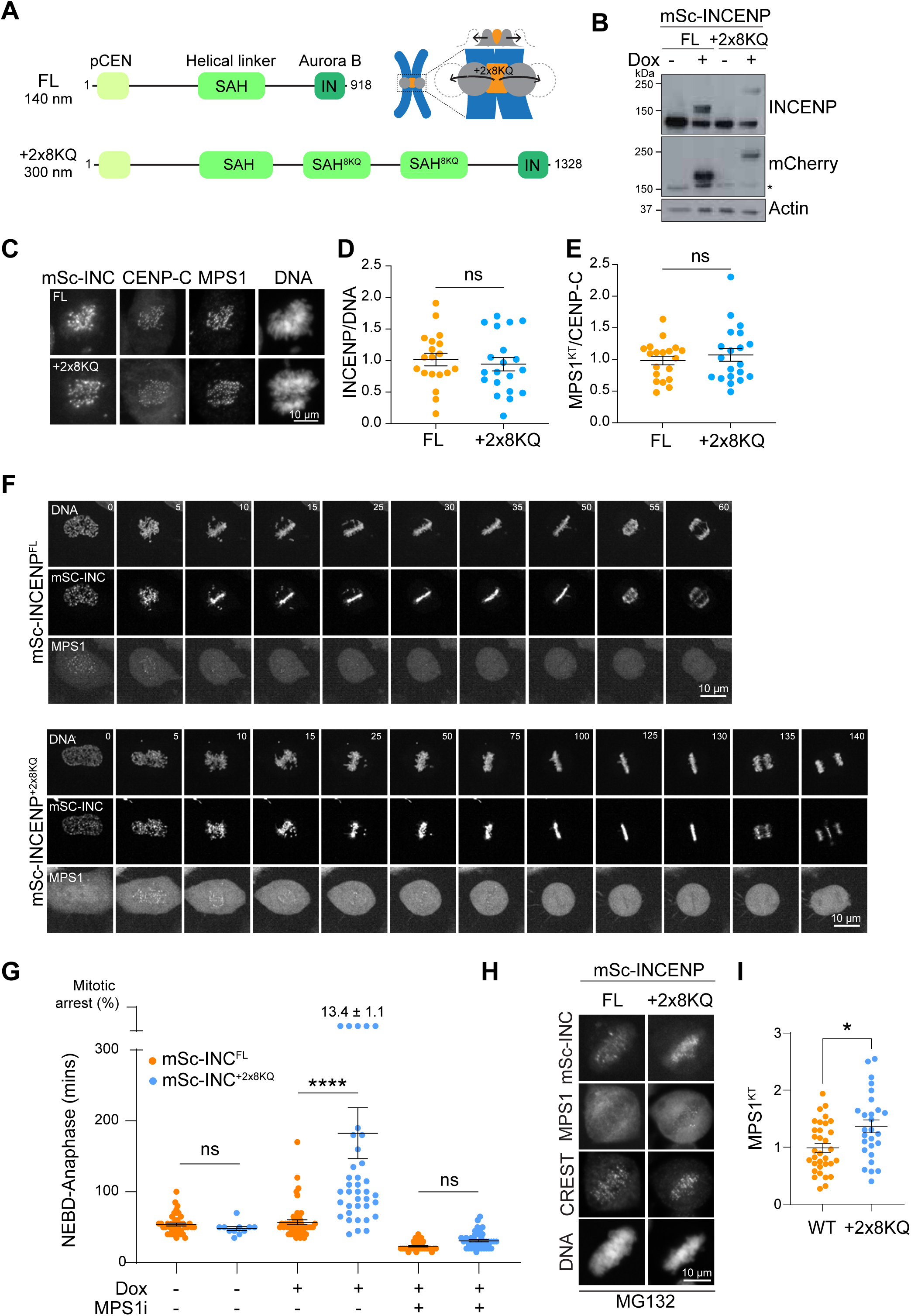
Extending INCENP prolongs outer-kinetochore Aurora B signalling and delays SAC silencing. (A) Schematic of the extended mScarlet-tagged INCENP. +2x8KQ indicates the addition of two SAH domains containing mutations of eight lysine residues to glutamine to impair microtubule binding (van der Horst et al., 2015). (B) HeLa Flp-In TREx cells expressing the indicated INCENP transgenes were lysed and blotted with antibodies against INCENP or mCherry (cross-reacts with mScarlet) with actin as a loading control. (C) HeLa Flp-In TRex MPS1-GFP prometaphase cells expressing mScarlet-INCENP^FL^ or mScarlet-INCENP^+2x8KQ^ were fixed and stained with antibodies against CENP-C. DNA was visualised with Hoechst. (D) Quantitation of pericentromeric localisation of INCENP and (E) MPS1 at kinetochores, as normalised to FL. Kruskal-Wallis statistical test was used where ns, p > 0.99. FL, n = 19; +2x8KQ, n = 20. (F) Stills for live imaging of the indicated cells going through unperturbed mitosis. Timings are in minutes. (G) Quantitation of mitotic timings (mean ± SEM), defined as nuclear envelope breakdown (NEBD) to anaphase, as shown in (F). FL, n = 43; +2x8KQ, n = 12; FL+Dox, n = 51; 2x8KQ+Dox, n = 38; FL+Dox+MPS1i, n = 32; FL+2x8KQ+Dox+MPS1i, n = 46. The graph shows the pooled data for at least 3 independent repeats. One-way ANOVA was used for statistical analysis where **** indicates p < 0.0001 and ns, p > 0.99. (H) HeLa Flp-In TREx cells were arrested in metaphase with MG132 and fixed for the analysis of MPS1-GFP kinetochore localisation. (I) Quantitation of (H) (mean ± SEM); FL, n = 31; +2x8KQ, n = 27. Kruskal-Wallis test was used for statistical analysis, where * is p <0.05.

Thus, extended versions of INCENP show mitotic delays due to a prolonged SAC. In contrast, truncation mutants of INCENP which retain pericentromeric localisation and Aurora B binding modules, but have a reduced distance between the two, are defective for MPS1 kinetochore targeting and fail to support the SAC. The most likely explanation for these observations is the altered spatial relationship between Aurora B and the outer kinetochore created by the greater or lesser reach of these INCENP variants from the pericentromeric chromatin. Taken together, the reciprocal effects of shortening and extending INCENP are consistent with INCENP length defining the effective range of Aurora B signalling at the outer kinetochore.

### NDC80 depletion impairs Aurora B-kinetochore separation while preserving SAC signalling

Microtubule-kinetochore attachment and MPS1 kinetochore recruitment are both reported to depend on the NDC80 complex (Cheeseman et al., 2006; DeLuca et al., 2006; Hiruma et al., 2015; Ji et al., 2015; Nijenhuis et al., 2013). To better understand the role of microtubule-dependent pulling forces we therefore explored how perturbation of the NDC80 complex affects these two processes. To achieve this, we used the Halo-PROTAC system which allows for rapid and specific protein destruction (Buckley et al., 2015) (Figure 4A).

**Figure 4.**
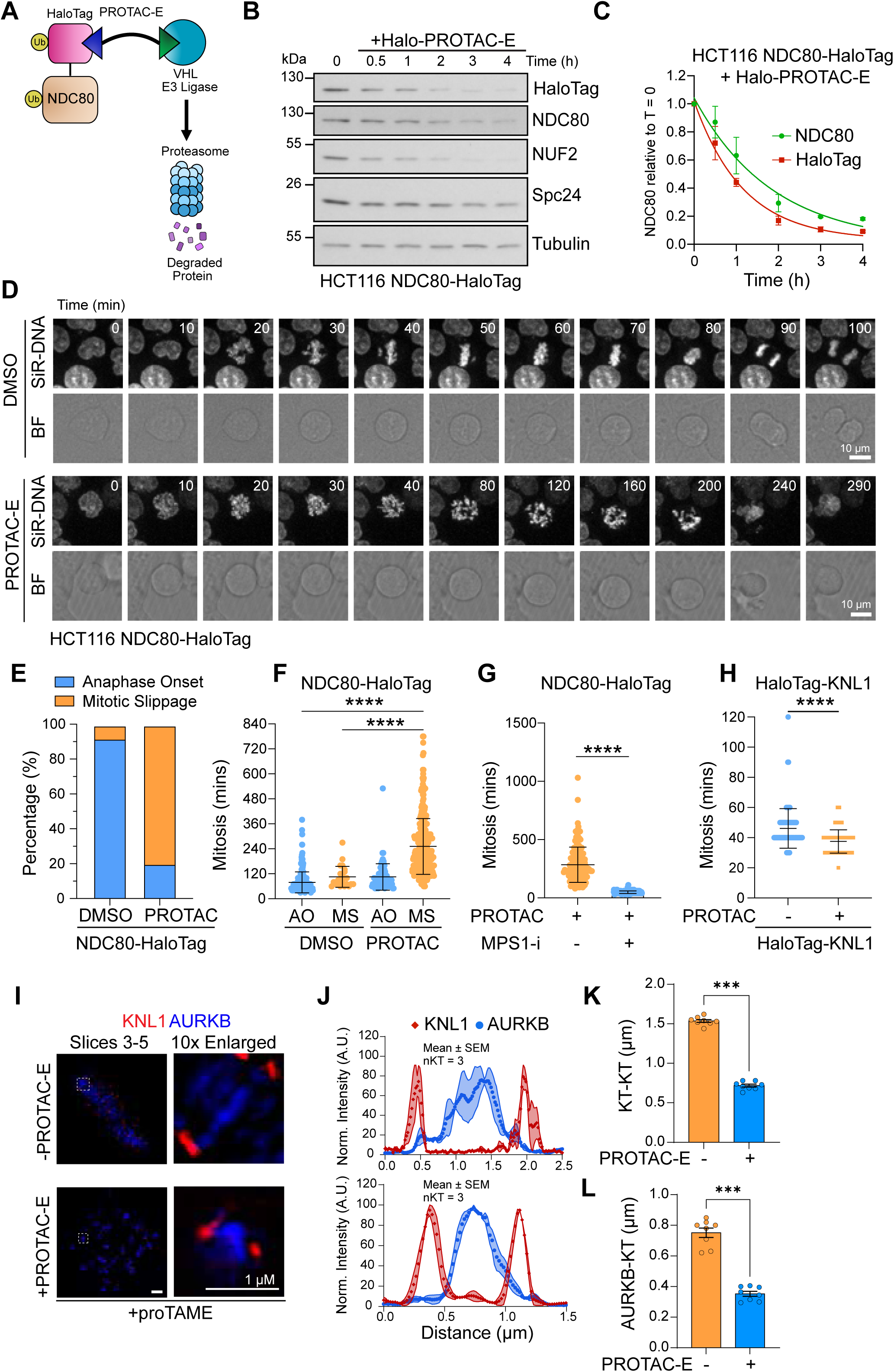
NDC80 depletion impairs Aurora B-kinetochore separation while preserving SAC signalling. (A) Basic schematic of Halo-PROTAC-E mediated proteolysis. (B) Halo-PROTAC-E degradation of asynchronous HCT116 NDC80-HaloTag cells, with samples taken over a 0–4-hour period. Sample protein concentrations were normalised by Bradford assay post lysis, and samples western blotted with the antibodies indicated. (C) Densitometry quantification of background-subtracted immunoblot bands for HaloTag and NDC80 over time (mean ± SD). Intensity values for each data set were normalised to T=0 (DMSO). One-phase decay curves were fitted to each dataset. (D) Control and PROTAC-E treated HCT116 NDC80-HaloTag cells were imaged at 10 min intervals progressing through mitosis. Chromatin was visualised with SiR-DNA; BF = bright field. HCT116 NDC80-HaloTag cells were treated for a minimum of three hours with Halo-PROTAC-E before the start of imaging. (E) Quantitation of the percentage of cells undergoing mitotic slippage in control and PROTAC-E treated cells from (D). (F) Mitotic timings in control and PROTAC-E-treated HCT116 NDC80-Halo cells (NEBD = Nuclear Envelope Breakdown, AO = anaphase onset; MS = mitotic slippage); mean ± SD; control AO, 78.7 ± 50.5 min, n = 262/286; control MS, 105.4 ± 50.3 min, n = 24/286; PROTAC AO, 99.6 ± 41.3 min, n = 79/403; PROTAC MS, 251.6 ± 134.1 min, n = 324/403. Two-tailed unpaired t test with Welch’s correction was used, **** p < 0.0001. (G) Mitotic timings in PROTAC-E treated cells ± MPS1 inhibitor (mean ± SD; control, 284.7 ± 151.1 min, n = 143; + MPS1-i, 49.5 ± 11.3 min, n = 157). (H) Mitotic timings in control and PROTAC-E-treated HCT116 KNL1-HaloTag cells (mean ± SD). For statistical analysis an unpaired t test with Welch’s correction was used. DMSO, n = 81; +PROTAC-E, n = 217. (I) Control and PROTAC-E treated HCT116 NDC80-HaloTag cells were arrested in mitosis with APC/C inhibitor proTAME and then fixed and processed for TauSTED imaging. Aurora B and outer kinetochore protein KNL1 were visualised by antibody staining. Representative maximum intensity projections of selected slices of kinetochore pairs are shown. The indicated kinetochore pairs are shown 10x enlarged at the side. (J) Line scans show signal intensity across kinetochore pairs (mean ± SEM, n = 3). (K) KT-KT (KNL1-KNL1) distances between kinetochore pairs from (I) are plotted (mean ± SD; **** indicates p < 0.0001). (L) Aurora B-KT (KNL1) distances are plotted (mean ± SEM; **** indicates p < 0.0001). Two-tailed, unpaired t-tests were used for statistical analysis.

When NDC80-Halo cells were treated with Halo-PROTAC-E, NDC80 and its partner NUF2 were reduced to less than 10% of starting levels within 4 h (Figure 4B and C). SPC24, another component of the outer kinetochore, was only partially reduced (Figure 4B). Cells treated for 3 h with Halo-PROTAC-E, then imaged passing through mitosis, failed chromosome alignment and did not form a metaphase plate (Figure 4D). Mitotic duration was markedly prolonged and 80.2% of cells eventually exited by mitotic slippage rather than anaphase (Figure 4D-F). The high proportion of mitotic slippage shows that chromosome segregation was failing, consistent with the idea that amphitelic microtubule attachments to the NDC80 complex generate the pulling forces that separate sister chromatids in anaphase in addition to their role in chromosome biorientation. The lengthened mitosis that we observed after NDC80 degradation was spindle checkpoint dependent as MPS1 inhibition shortened mitosis from 284.7 ± 151.1 min to 49.5 ± 11.3 min (Figure 4G). This suggests that despite a level of NDC80 depletion which abrogated normal force generation, residual NDC80 was sufficient to mediate a level of MPS1 recruitment which allowed SAC establishment and prolongation of mitosis. Of note, this phenotype was specific to NDC80 as degradation of KNL1 in KNL1-HaloTag cells resulted in shortened mitotic timing in comparison to control cells, most likely caused by the loss of the KNL1 molecular platform for MCC assembly (Musacchio, 2015) (Figure 4H and Figure S3A-C).

Amphitelic microtubule attachments to the NDC80 complex generate the pulling forces that separate pericentromeric chromatin from the outer kinetochore (Liu et al., 2009; Ruza et al., 2025). We therefore hypothesized that reduction in NDC80 levels was preventing separation of the outer kinetochore and pericentromere. Accordingly, high resolution TauSTED microscopy showed that NDC80 depletion reduced both kinetochore-kinetochore distance and the separation between Aurora B and KNL1 (Figure 4I-L). These distances were measured using the outer kinetochore checkpoint scaffold protein KNL1, since NDC80 was strongly reduced after PROTAC-E treatment.

Together, these data suggest that chromosome alignment and segregation, and checkpoint signalling have different functional thresholds for NDC80. Following PROTAC treatment there is too little NDC80 to support force generation and chromosome alignment, yet enough to support an MPS1-dependent SAC arrest. This difference provides an explanation for how reduced levels of NDC80 depletion can preserve checkpoint signalling while preventing the mechanical transition required for SAC silencing. Similar prolonged mitosis followed by slippage has been reported after auxin-induced NDC80 complex depletion (Kim et al., 2024). Taken together, these data support the conclusion that the SAC remains active when the microtubule-dependent pulling forces necessary for the separation of pericentromeric chromatin from the outer kinetochore cannot be established.

### Recruitment of Aurora B to bioriented outer kinetochores re-instates the SAC

Different mechanisms have been proposed to explain how kinetochore-microtubule attachment and tension are translated into SAC silencing during chromosome biorientation (Khodjakov and Pines, 2010; Nezi and Musacchio, 2009; Pinsky and Biggins, 2005). Tension generated by amphitelic kinetochore–microtubule attachments has been suggested to explain force-dependent attachment stabilisation and termination of error correction, and hence indirectly promotes checkpoint inactivation (Lampson and Cheeseman, 2011; Liu et al., 2009). In this model, tension is detected by the spatial separation of centromeric Aurora B from outer-kinetochore substrates. Alternatively, direct competition of MPS1 and microtubules for binding sites on the NDC80 complex was proposed to provide an attachment sensor for the SAC (Hiruma et al., 2015; Ji et al., 2015). However, it has remained unclear how these models relate to the essential role of Aurora B in promoting MPS1 recruitment and checkpoint establishment (Hayward et al., 2019b; Hayward et al., 2022; Nijenhuis et al., 2013; Santaguida et al., 2011; Saurin et al., 2011). A key observation in this regard is that MPS1 can localise to microtubule end-on attached kinetochores, and further findings that show MPS1 recruitment is Aurora B dependent (Hayward et al., 2022). The spatial separation model that we are testing here, makes a specific prediction in this regard. Force-dependent separation of Aurora B from the outer kinetochore during chromosome biorientation attenuates MPS1 recruitment. Therefore, artificially placing Aurora B close to the presumed MPS1 binding site at the outer kinetochore after biorientation has completed, should restore MPS1 recruitment despite continued microtubule-attachment and tension.

To test this idea, we adapted our rapamycin-dimerization system (Hayward et al., 2022), to recruit an mCherry-FRB-tagged INCENP^749-918^ fragment, containing the Aurora B-binding IN-Box, to Mis12-Myc-5xFKBP (Figure 5A, Figure S4A). Rapamycin induced rapid MPS1 recruitment to kinetochores in metaphase-arrested HeLa MPS1-GFP cells, reaching levels comparable to or greater than in nocodazole-treated cells within 2 min (Figure 5B and C). At this time, kinetochore-microtubule attachments remained cold stable, Astrin and Ska3 remained at kinetochores, and inter-kinetochore distance was unchanged (Figure 5D-F, Figure S4B-E). Recruited MPS1 underwent T-loop autophosphorylation indicative of activation (Mattison et al., 2007), while KNL1 MELT phosphorylation and MAD1 recruitment increased rapidly (Figure 5G-I, Figure S4D-G). These downstream responses required MPS1 activity and were sensitive to Aurora B inhibition. Thus, acute recruitment of Aurora B to the outer kinetochore is sufficient to restore MPS1 recruitment and SAC signalling in the continued presence of stable kinetochore-microtubule attachment and substantial inter-kinetochore stretch. SAC reactivation therefore precedes detectable attachment destabilisation, showing that microtubule attachment is not sufficient to exclude MPS1 from the outer kinetochore.

**Figure 5.**
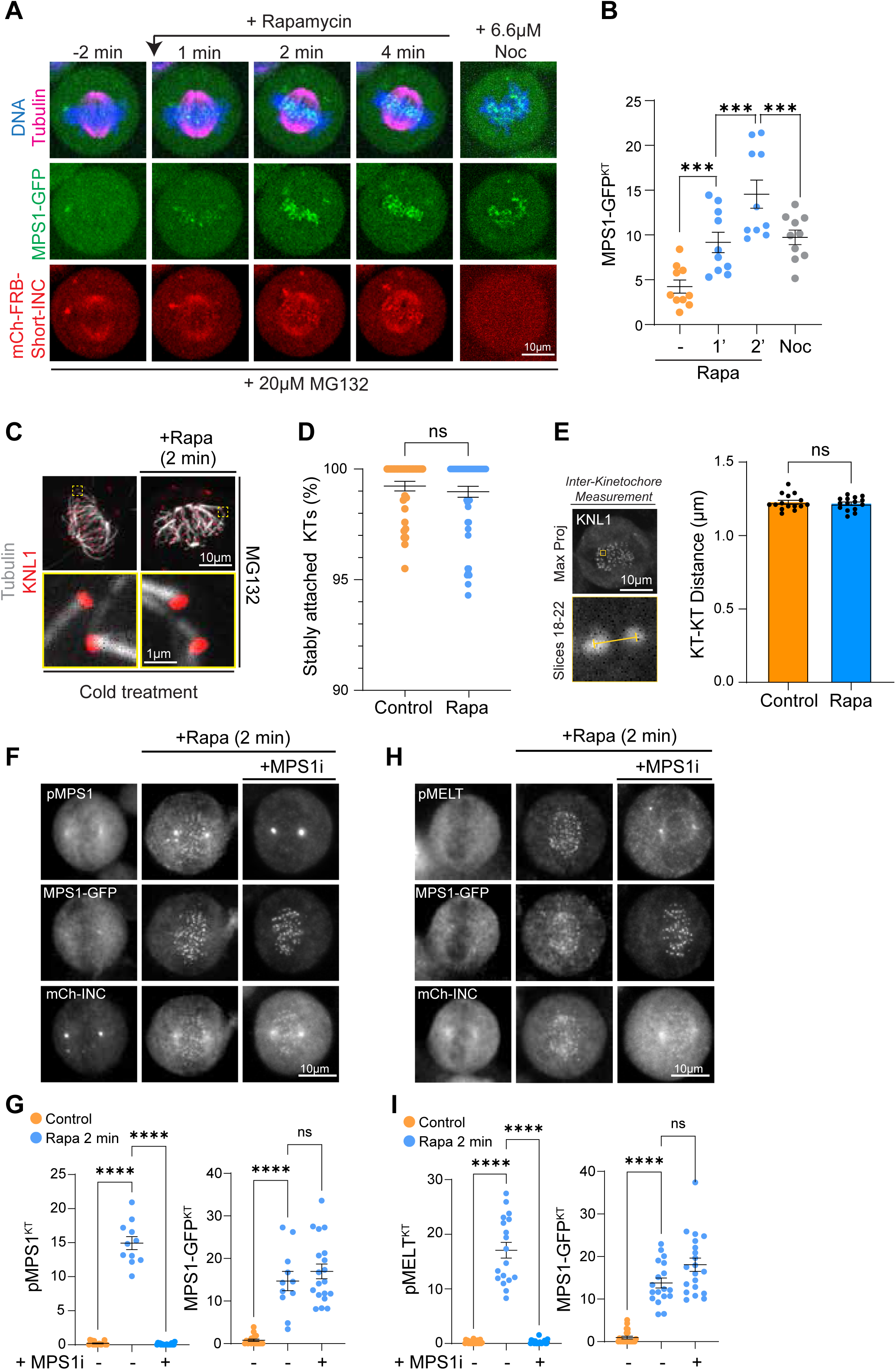
Recruiting Aurora B to microtubule-attached outer kinetochores re-engages SAC signalling. (A) Representative stills from live-cell imaging of HeLa MPS1-GFP cells expressing the IN-box-Aurora B kinetochore-targeting system. Cells were arrested in metaphase with MG132, treated with rapamycin, and imaged at 2 min intervals. (B) Quantification of kinetochore MPS1-GFP intensities in cells shown in (A), compared with nocodazole-arrested control cells. Graph shows mean ± SEM. *** indicates p < 0.001. n = 10 for all conditions. (C) Representative images from a cold-stability assay. Cells were arrested with MG132 and treated with or without rapamycin for 2 min before cold treatment and fixation. Cells were stained for α-tubulin and KNL1. (D) Quantification of cold-stable kinetochore–microtubule attachments in the conditions shown in (C). Graph shows mean ± SEM, Control, n = 41; +Rapa, n = 56. (E) Quantification of inter-kinetochore distances (KNL1-KNL1) in metaphase-arrested cells treated with or without rapamycin for 2 min. Control, n = 15; +Rapa, n = 15. (F) HeLa MPS1-GFP cells expressing the IN-box-Aurora B kinetochore-targeting system were arrested with MG132 and treated with rapamycin for 2 min, with or without the MPS1 inhibitor AZ3146. Cells were stained for MPS1-pT676 (MPS1-T-loop; pMPS1). (G) Quantification of kinetochore pMPS1 and MPS1-GFP in (F). Each dot represents a cell. Control, n = 18; +Rapa, n = 11; +Rapa +MPS1i, n = 17. Graph shows mean ± SEM. ****indicates p < 0.0001. (H) Cells were treated like in (F) and stained for phospho-MELT (pMELT). (I) Quantification of kinetochore pMELT and MPS1-GFP in cells in (H). Each dot represents a cell. Control, n = 22; +Rapa, n = 18; +Rapa + MPS1i, n = 20. Graph shows mean ± SEM. **** indicates p < 0.0001. For statistical analysis One-way ANOVA followed by Tukey’s multiple comparisons test was used.

The data presented here suggest that error correction and MPS1-dependent SAC signalling share a common upstream spatial-sensing principle. In the error-correction model, low-force attachments remain within the effective range of centromeric Aurora B and are destabilised by phosphorylation of outer-kinetochore substrates. Biorientation generates pulling forces that move these substrates away from Aurora B, reduce phosphorylation, and stabilise kinetochore-microtubule attachments (Lampson and Cheeseman, 2011; Liu et al., 2009). Repositioning Aurora B towards kinetochores resulted in constitutive error correction and checkpoint activity (Liu et al., 2009). Our results provide a molecular explanation for the latter observation by showing that Aurora B-outer kinetochore distance also directly regulates recruitment of MPS1, the kinase that initiates the checkpoint signal.

We therefore propose that SAC status is determined by the same upstream geometrical input as error correction. At low-force kinetochores, Aurora B remains within range of outer-kinetochore substrates, promoting attachment turnover and an outer-kinetochore state that supports MPS1 recruitment. MPS1 then generates the checkpoint signal and furthermore cooperates with Aurora B to mediate attachment error correction (Hayward et al., 2022). As correct bipolar attachments mature, microtubule-dependent pulling forces separate the outer kinetochore from pericentromeric Aurora B, favouring attachment stabilisation while limiting MPS1 recruitment. The downstream consequences are distinct; reduced phosphorylation of attachment-regulatory substrates for error correction and reduced MPS1-dependent checkpoint signalling for the SAC. Crucially, both are coupled to chromosome biorientation through the same spatial transition.

These events sit within a wider set of conditions required for checkpoint competence in mitosis. During mitosis, CDK1-CCNB1 licenses a spindle checkpoint-permissive mitotic state through regulation of both MPS1 and the Aurora B chromosome passenger complex (Alfonso-Perez et al., 2019; Hayward et al., 2019a). During this time window, PP2A-B56 counteracts Aurora B and MPS1 signalling at kinetochores to create dynamic cycles of checkpoint activation and silencing coupled to microtubule attachment (Foley et al., 2011; Hayward et al., 2019b; Hayward et al., 2022; Kruse et al., 2013; Suijkerbuijk et al., 2012; Xu et al., 2013). Our demonstration that Aurora B-outer kinetochore distance is a mechanical input that explains whether an attached kinetochore can either recruit or exclude MPS1 depending on its geometry will aid the identification of the crucial phosphorylated substrates and sites uniquely required for error correction, or MPS1 recruitment and checkpoint signalling. Our results are consistent with Aurora B-dependent phosphorylation of an outer-kinetochore component creating or exposing an MPS1-binding state.

An important focus for future work will therefore be to identify the Aurora B-regulated substrates that permit MPS1 recruitment, and to understand the mechanism by which they act. Previous work argues against a simple model in which phosphorylation of the NDC80 N-tail alone accounts for MPS1 recruitment (Hayward et al., 2022; Nijenhuis et al., 2013). Identifying the relevant Aurora B-regulated MPS1 recruitment sites is therefore important for understanding how microtubule-attachment is integrated with the kinase and phosphatase networks that regulate chromosome biorientation and checkpoint signalling.

## Acknowledgments

We thank our colleagues for discussion and advice, and the Oxford Leica Microsystems Centre of Excellence at the Micron Bioimaging Facility for their support & assistance in this work.

## Author contributions

Conceptualization: UG, FAB; Methodology: CWC, DBHG, DG, ML; Investigation: CWC, DBHG, DG, JD; Funding acquisition: FAB, UG; Supervision: FAB, UG; Writing – original draft: UG; Writing – review and editing: DBHG, CWC, DG, JD, FAB, UG.

## Methods

### Cell culture

Cell lines are validated stocks purchased from the ATCC. HeLa (#CRL-2.2) and derivatives were cultured in DMEM containing 1% [v/v] GlutaMAX™ (Gibco #10569010) and 10% (v/v) FBS. HCT116 cells (#CCL-247) were cultured in McCoy’s 5A medium (Gibco #16600082), supplemented with 10 mM sodium pyruvate (Thermo Fisher #11360088) and 10% (v/v) FBS. For routine passaging, cells were washed in PBS and then incubated with TrypLE™ Express Enzyme cell dissociation reagent (Gibco #12605036) for 5 min at 37°C, before resuspending detached cells in full medium for passage. All cell lines were maintained at 37°C, humidified 5% CO_2_ in a cell culture incubator (Thermo, HERAcell, #51013568). Mycoplasma negative status of cell lines was confirmed using the EZ-PCR Mycoplasma Test Kit with internal control (K1-0210, Geneflow). The cell lines used in our studies are not on the list as commonly misidentified lines.

### Generation of HCT116 NDC80-HaloTag MPS1-StayGold and HCT116 HaloTag-KNL1 cell lines

HCT116 NDC80-HaloTag cells have been described (Ruza et al., 2025). To generate HCT116 NDC80-HaloTag MPS1-StayGold cells, a guide RNA sequence 5’-AGTCCAATATATTTTATAGG -3’ targeting the C-terminal coding exon of MPS1 was inserted into a pX459 vector and delivered together with a donor homology template encoding a 12aa [GGSGGGGSGGGG]-linker-mStayGold-P2A-Neomycin resistance cassette, flanked by 600 bp homology arms on either side, by transient transfection. For transfection, transfection mixes were prepared in sterile DNase-free Eppendorf tubes (Eppendorf #0030108035) consisting of 100 µl Opti-MEM (Gibco #11058021), 3 µl Mirus TransIT-X2 transfection reagent (Mirus #MIR6004), and up to 1 µg of plasmid DNA. Transfection mixes were vortexed briefly for 20 s, and incubated at room temperature for 30 min, before adding dropwise to cells. Genomic DNA was isolated from clones using QuickExtract^TM^ DNA isolation solution (Lucigen #QE09050). PCRs were performed with KOD hot-start polymerase (Merck #71086), using primers flanking the gRNA recognition site. PCR products were ligated into pSC-A vectors using a Blunt-end PCR cloning kit (Agilent #240207). Ligated plasmids were transformed into XL-1 blue competent cells, and 10 colonies were mini-prepped (Qiagen #27107) and sent for Sanger DNA sequencing (Source Genomics, Cambridge, UK) to confirm the correct sequence for the edited alleles.

To generate HCT116 HaloTag-KNL1 cells, a guide RNA sequence 5’-AATGGATGGGGTGTCTTCAG-3’ targeting the N-terminal coding exon of KNL1 was inserted into a pX459 vector and delivered together with a donor homology template encoding Puro-Resistance-GSG-P2A-HaloTag-12aa [GGSGGGGSGGGG]-linker cassette flanked by 1000 bp homology arms on either side, by transient transfection.

### Generation of cell lines for rapamycin-mediated kinetochore recruitment of mCherry-FRB-IN-box

IN-box kinetochore recruitment by rapamycin addition was achieved by editing a previously described system (Ballister et al., 2014; Hayward et al., 2022), in which pERB109, Addgene #58280 had been adapted by placing miRFKBP5_Mis12-GFP-FKBP3 under doxycycline-inducible expression and replacing GFP with a Myc tag. Using Gibson assembly, the coding sequence for amino acids 47-748 was replaced from INCENP 47-918, which had been inserted after mCh-FRB. 831 and 804 bp homology arms flanking the AAVS1 safe harbour locus were also added, and the system was stably integrated into the AAVS1 safe harbour locus of HeLa cells homozygously expressing MPS1 endogenously tagged with GFP at the C-terminus by CRISPR/Cas9 knock-in, using the guide RNA sequence 5’-GTTAATGTGGCTCTGGTTCT-3’ (Addgene constructs #72833 and #72834)(Natsume et al., 2016).

### Generation of HeLa cell lines for MPS1 recruitment assays

HeLa Flp-In TREx cells engineered to express C-terminally GFP-tagged MPS1 have been described before (Ruza et al., 2025). Single integrated copies of the desired CPC transgene were introduced into HeLa Flp-In TREx GFP-MPS1 cells using the T-Rex doxycycline-inducible Flp-In system (Invitrogen). The pcDNA5/FRT/TO vector was used to make truncated or extended INCENP constructs N-terminally tagged with mScarlet. The truncated INCENP^Δ137-789^ was created by deletion mutagenesis; the 2xSAH^8KQ^ extension of INCENP was gene synthesised (GeneArt, ThermoFisher). Antibiotic-resistant clones were selected with 4 µg/ml blasticidin and 200 µg/ml hygromycin B, and successful modification was confirmed by western blotting.

### Reagents and antibodies

General laboratory reagents were obtained from Sigma-Aldrich and ThermoFisher Scientific unless specifically indicated. Inhibitors were obtained from Tocris Bioscience (PP1 and PP2A inhibitor Calyculin A (1336)); phosphatase inhibitor Okadaic acid (1136); MPS1 inhibitor AZ3146 (3994); APC/C inhibitor proTAME (7734/1)), Santa Cruz Biotechnology (Proteasome inhibitor MG132 (sc-201270)), Millipore (Thymidine (6060-5MG); Aurora B inhibitor ZM447439 (189410-5MG)), and Merck Chemicals Ltd (microtubule polymerization inhibitor Nocodazole (484728-10MG)), RO3306 (Insight – sc-358700). AZ3146 was used at 2 µM for 15 min before fixation for fixed cell analysis or added to cells 30 min before imaging for live cell imaging. ZM447439 was used at 10 µM for 5 min for fixed cell analysis. MG132 was added to cells 15-30 min before fixation to prevent exit from mitosis. proTAME was used at 20 µM for 2 hours to arrest cells in metaphase. Transfection reagents oligofectamine (Invitrogen) for siRNA oligos, and LT1 (Mirus-Bio) for plasmids were used according to the manufacturers’ instructions. Opti-MEM medium (Gibco) was used to set up transfections.

Commercially available antibodies were used for alpha-tubulin (Ms, mAb, DM1A Sigma-Aldrich T6199,), Aurora B/AIM1 (Ms mAb, BD Transduction, 611082, 1:1000 for western blotting, 1:2000 for immunofluorescence, 1:100 for TauSTED or Rb pAb Abcam ab2254 1:100 for 3D SIM); Borealin (Ms mAb, Santa Cruz, sc376635, 1:1000 for western blotting), CENP-C (guinea pig pAb; MBL (PD030); 1:2000 for immunofluorescence), CREST (Hu, 15-234, 1:2000 for immunofluorescence), beta-Actin (Ms mAb AC-15 HRP conjugate, abcam, ab49900; 1:5000 dilution for western blotting), HaloTag (Ms, mAb, Promega G9211 1:1000 for western blotting), Histone H3 (Rb pAb, Cell Signalling Technologies, 4499S, 1:5000 for western blotting), Histone H3 phospho-Serine 10 (H3pS10, Ms mAb, Cell Signalling Technologies, 9706S, 1:40000 for immunofluorescence), mCherry (Rb pAb, abcam, ab9106, 1:2000 for western blotting), MPS1 (mouse mAb (N1), abcam, ab11108, 1:2000 for western blotting), MAD1 (Rb pAb GeneTex GTX105079, 1;1000 for immunofluorescence), NDC80 (Ms, mAb, clone 9G3 Abcam ab3613, 1:1000 for western blotting), NUF2 (Ms, mAb, Clone E-6 Santa Cruz sc-271251 1:200 for western blotting), pan-AuroraA/B/C (Rb pAb, Cell Signalling Technologies, 2914S, 1:1000 for western blotting), SPC24 (Rb, pAb, ProteinTech 26268-1-AP, 1:1000 for western blotting), Survivin (Rb pAb, abcam, ab76424, 1:5000 for western blotting). Sheep-anti-phospho-KNL1 (pT875), sheep-anti phospho-MPS1 (pT676), sheep-anti-KNL1 and sheep antibodies against mCherry have been described (Bastos and Barr, 2010; Espert et al., 2014; Hayward et al., 2019b). For TauSTED/ 3D SIM, JFX646 or JFX554 were used to label NDC80-Halo cells respectively at 1:10,000. For SIM, donkey-anti-rabbit-DyLight-405 secondary antibody was used at 1:500 for Aurora B staining. For TauSTED, donkey-anti-mouse-AF594 was used at 1:250 for Aurora B staining. All other secondary antibodies were used at 1:1000 dilutions based on recommended stock concentrations. Secondary donkey antibodies against mouse, rabbit or sheep and labelled with Alexa Fluor 350, Alexa Fluor 555, Alexa Fluor 647 were purchased from ThermoFisher Scientific. Secondary donkey antibodies against guinea pig labelled with Alexa Fluor 647 and secondary donkey antibodies against mouse, rabbit or sheep labelled with HRP were purchased from Jackson ImmunoResearch Laboratories. Protein G conjugated to HRP were purchased from Merck. DNA dye Hoechst 33342 was purchased from ThermoFisher Scientific and used at a final concentration of 5 µg/ml.

For western blotting, proteins were separated by SDS-PAGE and transferred to nitrocellulose using a Trans-blot Turbo system (Bio-Rad). Protein concentrations were measured by Bradford assay using Protein Assay Dye Reagent Concentrate (Bio-Rad). All western blots were revealed using ECL (GE Healthcare).

### TauSTED and SIM Sample Preparation using HCT116 NDC80-HaloTag MPS1-mStayGold cell line

Cells were seeded onto high-precision 22 mm × 22 mm #1.5H coverslips (Marienfeld #0107052). To arrest cells in G2, 6 µM of the Cdk1-inhibitor RO-3306 (Tocris #4181) was added 18 hours prior to fixation. RO-3306 was washed out 4x in media containing the proteasome inhibitor MG132 (Sigma-Aldrich #474790) to trap prometaphase cells for 1-2 hours. Either 100 nM JFX-646 or JFX-554-HaloTag dye was added 15 minutes prior to fixation to fluorescently label NDC80-HaloTag (Grimm et al., 2017). For TauSTED, cells were then fixed for 12 minutes with G-PTEMF (0.1% [vol/vol] glutaraldehyde, 20 mM PIPES pH 6.8, 0.2 % [vol/vol] Triton X-100, 10 mM EGTA, 1 mM MgCl_2_, and 4% [vol/vol] formaldehyde). Fixation was quenched for 15 minutes with 50 mM NH_4_Cl. Coverslips were then incubated with primary antibodies for 1.5-2 hours, followed by a PBS wash and a further 1.5-hour incubation with secondary antibodies. Cells were then post-fixed for 15 minutes with 0.1% [vol/vol] glutaraldehyde and 3 % [vol/vol] formaldehyde to stabilise bound antibodies. Fixation was quenched with 1 mg/ml sodium borohydride for 7 minutes, followed by 100 mM glycine for 15 minutes. Coverslips were mounted onto cover glass using Everbrite mounting medium (Biotium #BG23001) and sealed with nail varnish. Samples were imaged the next day on a Leica STELLARIS TauSTED Xtend system, equipped with an NKT-Photonics 470-790 nm White Light Laser (WLL) for excitation, and MPB-589 nm and MPB-775 nm depletion lasers. Imaging was performed using the system’s CFL 100x/1.4NA Oil objective and HyD X detector, with acquisition controlled by Leica’s LAS X 4.8 software. Z-stacks were recorded with an average z-spacing of 0.3–0.4 µm.

For 3D-SIM samples with MPS1-mStayGold integration, the fixation method was modified as follows: Cells were fixed for 12 minutes with PTEMF (20 mM PIPES-KOH pH 6.8, 0.2% [vol/vol] Triton X-100, 10 mM EGTA, 1 mM MgCl_2_, and 4% [vol/vol] formaldehyde) without quenching. Antibody incubation and PBS washes were performed as described above for TauSTED samples. Coverslips were mounted onto microscope slides using Mowiol and left to cure overnight. 3D-SIM imaging was performed on a DeltaVision OMX SR system (GE Healthcare; IMSOL) equipped with a 60x/1.5 NA UPLAPO60XOHR oil-immersion objective (Olympus), pco.edge 4.2 sCMOS cameras (PCO), and 405, 488, 568, and 640 nm lasers. Data acquisition and reconstruction were performed as previously described (Ochs et al., 2024). Briefly, 15 raw images (5 phases × 3 angles) were captured at each z-position, and stacks were acquired at 125 nm z-steps. The objective correction collar was set to 0.130, an empirically determined value for imaging adherent cells (±4 μm), to minimise spherical aberration during reconstruction. Images were reconstructed in softWoRx 7.2.2 (GE Healthcare) using previously acquired, channel-specific optical transfer functions (OTFs) (Demmerle et al., 2017) and a Wiener filter setting of 0.0030, yielding a final reconstructed resolution of approximately 100-130 nm laterally (x-y, wavelength-dependent) and approximately 300 nm axially (z). SIMcheck (Ball et al., 2015) was used for quality control, and chromatic shift between camera channels was corrected using Chromagnon 0.94 3D alignment software (Matsuda et al., 2018) referencing same-day 3D-SIM acquisitions of multicolour EdU-labelled C127 epithelial cells as a colocalisation standard (Kraus et al., 2017).

### Image processing and data analysis for TauSTED and SIM images

Image analysis was performed using the Fiji distribution of ImageJ (V2.14). Quantification was performed on a per-kinetochore-pair basis (either on a single slice or a maximum projection of a limited number of slices). Line scans were generated by measuring the signal spanning the pericentric region between two kinetochore pairs. The same line was used to measure density across all channels. Pixel distances were converted to µm based on the scaling of each image at the time of acquisition. These data were then normalised in GraphPad Prism Version 10 for MacOS, with 0 defined as the smallest value in each data set and 100% as the largest mean in each data set. For SAC signals, this normalisation was performed manually relative to un-tensioned conditions, as tensioned kinetochores show no SAC signal. Normalising otherwise resulted in amplified noise. Each scan was plotted with ± SEM, shown as a lightly coloured region around the curve. Inter-kinetochore distances were measured using a 1-px-wide line in Fiji, with each end placed centrally on the kinetochore. Aurora B signal-to-kinetochore distances were determined by fitting a Gaussian curve to each signal and measuring the distance between the peaks. A single pair of kinetochores was treated as a replicate within a cell (n). Statistical analysis of these distances was performed using a parametric two-tailed unpaired t-test. Graphs display the mean ± SD, with p-values shown on the graphs as follows: p > 0.05 (not significant, ns), p < 0.05 (*), p < 0.01 (**), p < 0.001 (***), p < 0.0001 (****).

### Cold-stability assays

HeLa MPS1-GFP cells expressing the mCherry-FRB-IN-Box/Aurora B targeting system were seeded in 6-well plates at a density of 80,000 cells per well, with duplicate #1.5 thickness coverslips in each well. 24 h after seeding, cells were supplemented with 2 µM doxycycline and incubated for a further 48 h. Following doxycycline treatment, culture medium was aspirated and the plate was placed on ice for 9 min to induce depolymerisation of unstable microtubules (Rieder, 1981). During cold treatment, 2 mL of ice-cold culture medium (DMEM with 1% (vol/vol) GlutaMAX (Life Technologies) containing 10% (vol/vol) bovine calf serum) was added to each well to prevent the cells from drying. Following the 9 min incubation, the ice-cold medium was aspirated and cells were fixed at room temperature, adding 2 mL PTEMF to each well. Cells were subsequently processed according to the standard immunofluorescence staining protocol described above. Microtubules were visualised by staining for α-tubulin and kinetochores by staining with anti-KNL1 antibodies. The number of cold stable microtubules per cell was then manually determined as the fraction of microtubule-associated KNL1-foci of the total number of KNL1-foci per cell.

### Inter-kinetochore distance measurements

For inter-kinetochore (inter-KT) distance measurements presented in Figure 5, images were acquired using an Olympus SoRa spinning-disk confocal microscope equipped with a 60×/1.5 NA objective on an Olympus IX-83 microscope stand. Images were acquired using 3.2× optical zoom and the Yokogawa CSU-W1 SoRa super-resolution system to enable visualisation of individual kinetochore pairs. Where kinetochore pairs extended across multiple focal planes, the relevant z-stacks were projected to allow both kinetochores within a pair to be visualised simultaneously. Inter-KT distances were measured manually in Fiji by drawing a 1-pixel wide line from the centre of one kinetochore to the centre of its paired kinetochore. The resulting centre-to-centre distance was recorded for each kinetochore pair and used for subsequent comparison between experimental conditions.

### MPS1 recruitment assays

To test the role of the INCENP long, flexible arm in MPS1 recruitment, cells were depleted of endogenous INCENP by RNAi and fixed or arrested in mitosis and fixed. Cells were grown on coverslips (16 mm circular No. 1 ½, ThermoFisher) placed in 6 well plates a day before treatment. All siRNA depletions were performed for 48 h and all siRNAs were purchased from Dharmacon. Control siRNA was performed against luciferase as a control (5’-CGUACGCGGAAUACUUCGAUU-3’). siRNA targeting the 3’-UTR of INCENP was: INCENP (5’-GGCUUGGCCAGGUGUAUAUdTdT-3’). For rescue experiments, 2 µM Doxycycline was added 30 min before siRNA addition to induce mScarlet-transgene expression. A second Doxycycline induction was performed 24 hours into siRNA depletion.

To arrest the cells in a spindle checkpoint active prometaphase-like state and depolymerise microtubules, 0.66 µM nocodazole was added for 2 h. To prevent mitotic exit in cases where the spindle checkpoint was compromised, 20 µM of the proteasome inhibitor MG132 was added 30 min before fixation.

Cells were fixed with PTEMF buffer (20 mM PIPES-KOH pH 6.8, 0.2% [v/v] Triton X-100, 10 mM EGTA, 1 mM MgCl_2_, 4% [v/v] formaldehyde) for 12 minutes at room temperature. Coverslips were incubated in blocking buffer (3% [wt/v] bovine serum albumin) for a minimum of 1 hour. Coverslips were incubated face-down on 80 µl droplets of primary antibodies in a humidified chamber for 1 hour. Following primary antibody incubation coverslips were washed 3 x in PBS and incubated face-down on 80 µl droplets of diluted antibodies in a humidified chamber for 45 minutes. Coverslips were washed 3 x in PBS and 1 x in distilled deionised water. Coverslips were left to dry completely before being mounted with 7 µl of Mowiol 4-88 (Sigma) according to manufacturer’s instructions. Samples were imaged using a 60x/1.35-NA oil-immersion objective on a BX61 Olympus microscope equipped with filter sets for DAPI, EGFP/Alexa Fluor 488, 555 and 647 (Chroma Technology), a CoolSNAP HQ2 camera (Roper Scientific), and MetaMorph 7.5 imaging software (GE Healthcare). Each image consisted of a total stack of 2 µm with 0.2-µm intervals.

### Image processing and data analysis for MPS1 recruitment assays

Image processing was performed using the Fiji distribution of ImageJ. Quantification was performed on sum-projected images whilst figures shown were collated from maximum projections. Individual cells were cropped to 250 x 250-pixel images. Image analysis was performed on FIJI and MATLAB R2023b (MathWorks, Natick, MA, USA). Kinetochore intensities for each fluorescence channel were determined by segmentation of individual, non-overlapping kinetochores of at least 8 px in diameter (which defines the kinetochore ROI) and a measurement of the mean pixel intensity of each channel within the kinetochore ROI. A minimum of 20 kinetochores were measured per cell. Its corresponding background ROI was derived from the mean intensity in the nearest 52 pixels surrounding each kinetochore ROI, without encroaching into other kinetochore ROIs, using the k-nearest neighbours (KNN) algorithm. For each cells, kinetochore signal intensities were background-adjusted by subtracting the background signal on a channel-by-channel basis. For each kinetochore within a cell the intensity of the channel of interest was normalised by dividing by the intensity of the corresponding CENP-C intensity. Mean kinetochore localization intensities were then calculated for each cell.

### Statistical analysis for MPS1 recruitment assays

All statistical analysis was performed using GraphPad Prism Version 10 for Windows (GraphPad Software, San Diego, CA, USA). Each cell measured was considered as a biological replicate (n), hence mean measurements calculated for each cell were used for statistical analysis. At least three independent repeats of each experiment were performed, with statistical analysis performed using biological replicates from independent experiments. The precise n values, where n is the number of cells or kinetochore pairs analysed, are indicated in the figures or figure legends. If only two groups were compared an unpaired two-tailed t-test was used (with Welch’s correction if the groups had unequal standard deviations). If more than two groups were compared with equal standard deviations a one-way ANOVA was used. Graphs display the mean± SEM. p-values are shown on graphs as follows: p > 0.05 = not significant (ns), p < 0.05 = *, p < 0.01 = **, p < 0.001 = ***, p < 0.0001 = ****.

### Immunoprecipitations

Cells were grown in 15 cm dishes (number of dishes varied depending on the size of immunoprecipitation). mSc-INCENP Flp-In transgene expression was induced by addition of 2 μM doxycycline, in conjunction with depletion of endogenous INCENP, for 48 h. In all experiments, cells were arrested in 0.1 µM nocodazole for 18 h and harvested by mitotic shake-off. Cells were pelleted at 500 xg for 4 min at room temperature, washed with PBS and pelleted again. Pellets were lysed in lysis buffer (20 mM Tris-HCl pH 7.4, 150 mM NaCl, 1% (vol/vol) IGEPAL, 0.1% (wt/vol) sodium deoxycholate, 100 nM okadaic acid, 40 mM sodium βglycerophosphate, 10 mM NaF, 0.3 mM sodium vanadate, 1:250 protease inhibitor cocktail (Sigma), 1:100 phosphatase inhibitor cocktail (Sigma)) for 30 min at 4°C before being clarified at 14,000 rpm for 15 min at 4°C. Per 1 mg of lysate, 1 µg appropriate antibody was added (sheep anti-mCherry or sheep anti-GST as a negative IgG control). Immunoprecipitations were performed using Protein-G Dynabeads (Invitrogen). All beads were washed 3x in lysis buffer prior to use. Immunoprecipitations were performed at 4°C for 2 h. Beads were washed 6x, 3x in lysis buffer and 3x in wash buffer (20 mM Tris-HCl pH 7.4, 150 mM NaCl, 0.1% (vol/vol) IGEPAL, 40 mM sodium β-glycerophosphate, 10 mM NaF, 0.3 mM sodium vanadate). Beads were separated from the flow-through using a magnet. If samples were used for western blot analysis, 3x Laemmli buffer was added and samples denatured at 95°C for 5 min.

### Live cell imaging

For high-resolution live cell imaging of HeLa MPS1-GFP cells expressing the IN-box/Aurora B kinetochore targeting system and for live cell imaging of HeLa Flp-In-TRex MPS1-GFP cells, cells were seeded on circular glass bottom Fluorodish imaging dishes (World Precision Instruments) in Leibovitz’s L-15 media (ThermoFisher Scientific) supplemented with 20% FBS and imaged at 37°C with 5% CO_2_ on an Olympus SoRa spinning disk confocal microscope using a 60x/1.5-NA or 100x/1.45-NA objective fitted to an Olympus IX-83 microscope stand with a Yokogawa CSU-W1 SoRa super-resolution spinning disk. Solid state lasers emitting 405 nm, 488 nm, 561 nm and 633 nm were used. Images were captured with a Prime 95B sCMOS camera (photometrics) using the Olympus cellSens software package.

For Rapamycin experiments, 10 µm stacks with intervals of 0.5 µM of MG132- or nocodazole-arrested cells were imaged using the 60x/1.5-NA objective at intervals of 2 min over a total period of 22 minutes, with 100 µl PBS + Rapamycin added in the interval between the first and second timepoints. Microtubules and DNA were detected with 50 nM SiRTubulin (Spirochrome) and 4 µM Hoechst-33342 added 1 hour prior to imaging. Fluorescence intensities of kinetochore foci in live-cell imaging experiments were quantified in FIJI using circular regions of interest (ROIs), with background subtraction applied to all measurements. Unless otherwise stated, five 8-pixel-diameter circular ROIs were positioned over foci of interest, and fluorescence intensity was measured for each ROI. Background fluorescence was determined from five ROIs positioned in chromatin-free cytoplasmic regions and was subtracted from the corresponding foci measurements. For analysis of kinetochore MPS1-GFP and mCherry-FRB-IN-Box fluorescence during live-cell imaging, five 8-pixel-diameter circular ROIs were positioned over MPS1-positive kinetochores at the first timepoint at which MPS1-positive kinetochores were visible. The same ROIs were then used to measure fluorescence at preceding timepoints. At subsequent timepoints, ROIs were repositioned as required to track the same kinetochores throughout the time course. Where no MPS1-positive kinetochores were visible, five ROIs were instead positioned within the mitotic plate, as defined by Hoechst staining or outline of chromatin in other channels.

### Determination of mitotic timings

To determine cumulative mitotic exit timings of HeLa Flp-In TREx MPS1-GFP cell lines, cells were seeded on 6-well #1.5H glass-bottomed dishes (Cellvis) in Fluorobrite media (ThermoFisher Scientific) supplemented with 10% FBS and 1x GlutaMAX (ThermoFisher Scientific). Imaging was performed using a 20x/0.75 NA air objective on an EVOS M7000 (ThermoFisher Scientific) with software version 2.0.2094.0, equipped with an onstage incubator and DAPI, GFP, Texas Red, and Cy5 light cubes. Cells were synchronised in 2 mM Thymidine overnight before release by washing out into imaging media. SiRDNA (50 nM, Spirochrome was added 8 hours prior to imaging. Cells were imaged at 37°C under 5% CO_2_ every 5 minutes for 12 hours.

To determine mitotic timings of HCT116 NDC80-HaloTag cells, cells were seeded on 35-mm dishes, with a 14 mm 1.5 thickness cover glass bottom (MatTek #P35G-1.5-14-C) and grown for at least 2 days in McCoy’s 5A medium (Gibco #16600082), supplemented with 10 mM sodium pyruvate (Thermo Fisher #11360088) and 10% (v/v) FBS. They were then treated with 50 nM SiR-DNA (Spirochrome #Sc005), and either DMSO or 300 nM Halo-PROTAC-E for at least 3 hours. Dishes were then placed into an environment chamber (Tokai Hit) mounted onto the microscope stage to maintain cells in a 37°C and 5% CO_2_ environment throughout the imaging duration. Imaging was performed on an Ultraview Vox spinning disk confocal system running Volocity software (PerkinElmer, V6.3.0) using a 60x/1.42 NA UPlanSApo oil objective on an Olympus IX-81 inverted microscope equipped with an electron multiplying charge coupled device (EM-CCD) camera (Hamamatsu Photonics no. C9100-13). Dependent on the experiment, 488 nm, 561 nm, or 651 nm lasers were used with exposures ranging from 50-200 ms at 1.5-7% laser power; brightfield images at 30-50 ms were also taken for cell morphology reference images. Multi-point acquisition was performed, with images taken over 21-24 planes at 0.9 µm spacing; time intervals are specified in each figure. Maximum intensity projection, image cropping, time in mitosis durations, and intensity quantifications were all performed in Fiji.

