## Supplemental figures for "Spatial separation provides the force-coupling mechanism for the spindle assembly checkpoint"

#### Supplemental data

#### Supplemental Figure legends

**Figure S1. Cells expressing truncated INCENP have normal Aurora B activity.** (A) HeLa Flp-In TREx cells depleted of endogenous INCENP and expressing endogenously GFP-tagged MPS1 as well as the indicated INCENP transgenes were either synchronised with nocodazole (or left asynchronous (B)) and fixed and stained with the indicated antibodies. (C) and (D) Quantitation of histone H3 pSer10 in (A) and (B), respectively. siCon, n = 27; siINC, n = 29; FL, n = 32; ΔSAH, n = 27; Δ137-789, n = 36 for cells in nocodazole and siCon, n = 22; siINC, n = 29; FL, n = 25; ΔSAH, n = 21; Δ137-789, n = 25 for asynchronous cells. (E) HeLa Flp-In TREx cells as in (A) were mitotically arrested in 0.66 µM Nocodazole and 10 µM MG132 for 2 hours and 15 min, respectively, prior to fixation and staining with the indicated antibodies. (F) Quantitation of the mScarlet-INCENP signal in cells from (E), mean ± SEM. siCon, n = 15; siINC, n = 18; FL, n = 16; ΔSAH, n = 36; Δ137-789, n = 28 (G) Quantification of kinetochore (KT) MPS1 signal normalised to kinetochore CENP-C in cells in (E) siCon, n = 15; siINC, n = 18; FL, n = 16; ΔSAH, n = 37; Δ137-789, n = 25; (F) siCon, n = 27; siINC, n = 29; FL, n = 32; ΔSAH, n = 27; Δ137-789, n = 36 and (G) siCon, n = 22; siINC, n = 29; FL, n = 25; ΔSAH, n = 21; Δ137-789, n = 25. Kruskal-Wallis test was used for statistical analysis, where **** p<0.0001, *** p<0.001, ** p<0.01, * p<0.05. Scale bar, 10 µm.

**Figure S2. Mitotic progression in cells expressing truncated versions of INCENP.** (A), (B), (C) HeLa Flp-In TREx cells depleted of endogenous INCENP and expressing the indicated INCENP transgenes were synchronised in early S-phase with 2 mM thymidine and released for 8 hrs before unperturbed live imaging. Timings are in minutes. Note that cells expressing truncated versions of INCENP lacking the SAH are defective for INCENP localisation to the central spindle, as previously described (Vader et al., 2007). (D) Quantification of cells with chromosome alignment or spindle assembly checkpoint failure in (A)-(C). Mitotic defects plotted as averaged percentage from 3 biological repeats, where n>10 per repeat. (E) Quantification of mitotic timings. siCon, n = 31; siINC, n = 33; FL, n = 32; ΔSAH, n = 36; Δ137-789, n = 50. Graph shows mean ± SEM.

**Figure S3. Mitotic progression in PROTAC-E treated HCT116-HaloTag-KNL1 cells.** (A) Halo-PROTAC-E degradation of asynchronous HCT116 HaloTag-KNL1 cells, with samples taken over a 0–4-hour period. Sample protein concentrations were normalised by Bradford assay post lysis, and samples western blotted with the antibodies indicated. (B) Densitometry quantification of background-subtracted immunoblot bands for HaloTag and KNL1 over time (mean ± SD). Intensity values for each data set were normalised to T=0 (DMSO). One-phase decay curves were fitted to each dataset. (C) Control and PROTAC-E treated HCT116 HaloTag-KNL1 cells were imaged at 10 min intervals progressing through mitosis. Chromatin was visualised with SiR-DNA; BF = bright field. HCT116 HaloTag-KNL1 cells were treated for a minimum of three hours with Halo-PROTAC-E before the start of imaging.

**Figure S4. Rapamycin-mediated IN-box kinetochore recruitment leads to rapid re-initation of the SAC.** (A) i. Schematic of the rapamycin-dependent dimerization system used to recruit the shortened INCENP–Aurora B complex to the kinetochore protein MIS12. ii. Schematic of the experimental design aimed at bypassing the spatial separation of Aurora B and outer kinetochore. (B) mCherry-FRB-IN-box was precipitated from cells expressing the IN-box-rapamycin recruitment system in the presence or absence of rapamycin. The blot was probed with the indicated antibodies. (C) and (D) Cells expressing the IN-box-rapamycin recruitment system were treated with rapamycin for 2 min, fixed and stained with the indicated antibodies. (E) and (F) Quantifications of the imaging in (C) and (D). For (E), control, n = 36; +Rapa, n = 51. For (F), control, n = 25; +Rapa, n = 30. (G) Cells expressing the IN-box-rapamycin recruitment system were treated with rapamycin or DMSO for 2 min in the presence or absence of Aurora B inhibitor ZM447439 and stained for MAD1. (H) Quantification of MAD1 and MPS1-GFP in (G). Each dot represents a cell. Control, n = 25; +Rapa, n = 17; +Rapa +AurBi, n = 21. (I) Cells expressing the IN-box-rapamycin recruitment system were treated with rapamycin or DMSO for 2 min in the presence or absence of Aurora B inhibitor ZM447439 and stained for KNL1-pMELT. (J) Quantification of pMELT and MPS1-GFP in (I). Each dot represents a cell. Control, n = 22; +Rapa, n = 18; +Rapa +AurBi, n = 22. (K) Cells expressing the IN-box-rapamycin recruitment system were treated with rapamycin or DMSO for 2 min in the presence or absence of MPS1 inhibitor AZ3146 and stained for MAD1 and KNL1. (L) Quantification of MAD1 and MPS1-GFP in (K). Each dot represents a cell. Control, n = 25; +Rapa, n = 17; + Rapa + MPS1i, n = 22. Graph shows mean ± SEM. For statistical analysis, one-way ANOVA followed by Tukey's multiple comparisons test was used.

#### Figure S1

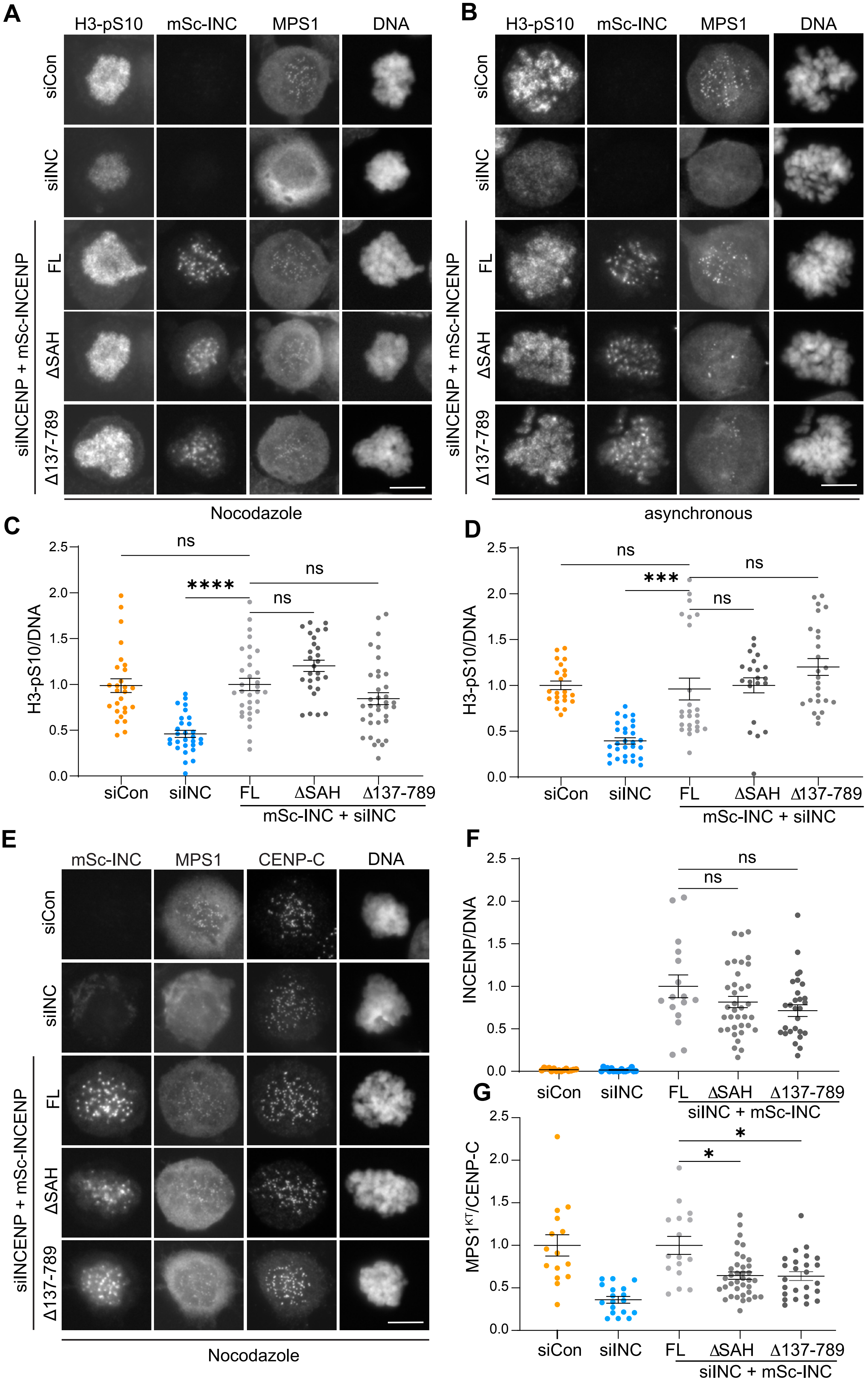
A
H3-pS10
mSc-INC
MPS1
DNA
siCon
siINC
siINCENP + mSc-INCENP
FL
ΔSAH
Δ137-789
Nocodazole
B
asynchronous
C
H3-pS10/DNA
ns
****
siCon
siINC
FL
ΔSAH
Δ137-789
mSc-INC + siINC
D
***
E
CENP-C
F
INCENP/DNA
siINC + mSc-INC
G
MPS1KT/CENP-C
*

### Figure S2

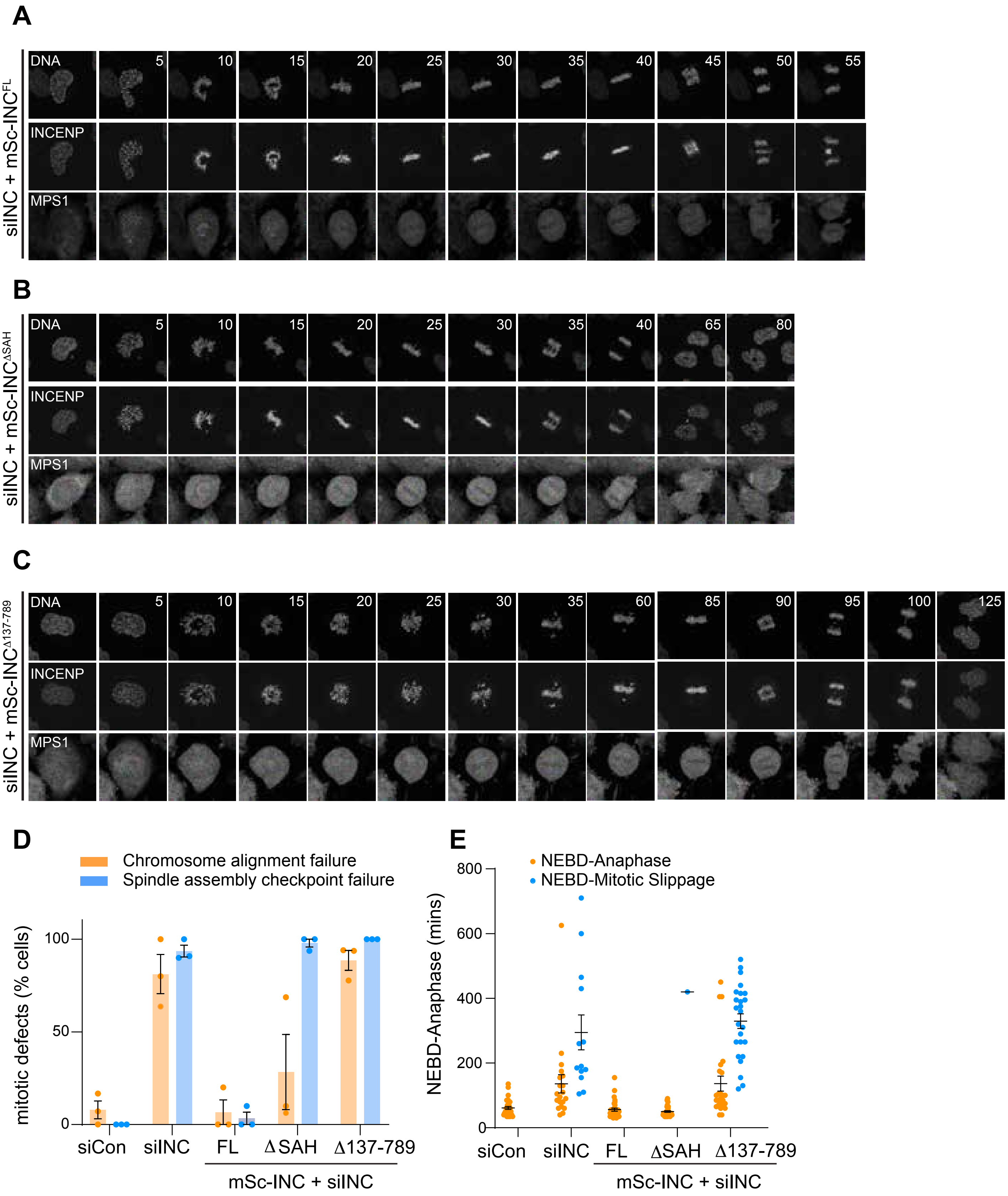
A
siINC + mSc-INC^FL
DNA
INCENP
MPS1
5 10 15 20 25 30 35 40 45 50 55
B
siINC + mSc-INC^ΔSAH
DNA
INCENP
MPS1
5 10 15 20 25 30 35 40 65 80
C
siINC + mSc-INC^Δ137-789
DNA
INCENP
MPS1
5 10 15 20 25 30 35 60 85 90 95 100 125
D
Chromosome alignment failure
Spindle assembly checkpoint failure
mitotic defects (% cells)
0
50
100
siCon
siINC
FL
ΔSAH
Δ137-789
mSc-INC + siINC
E
NEBD-Anaphase
NEBD-Mitotic Slippage
NEBD-Anaphase (mins)
0
200
400
600
800
siCon
siINC
FL
ΔSAH
Δ137-789
mSc-INC + siINC

### Figure S3

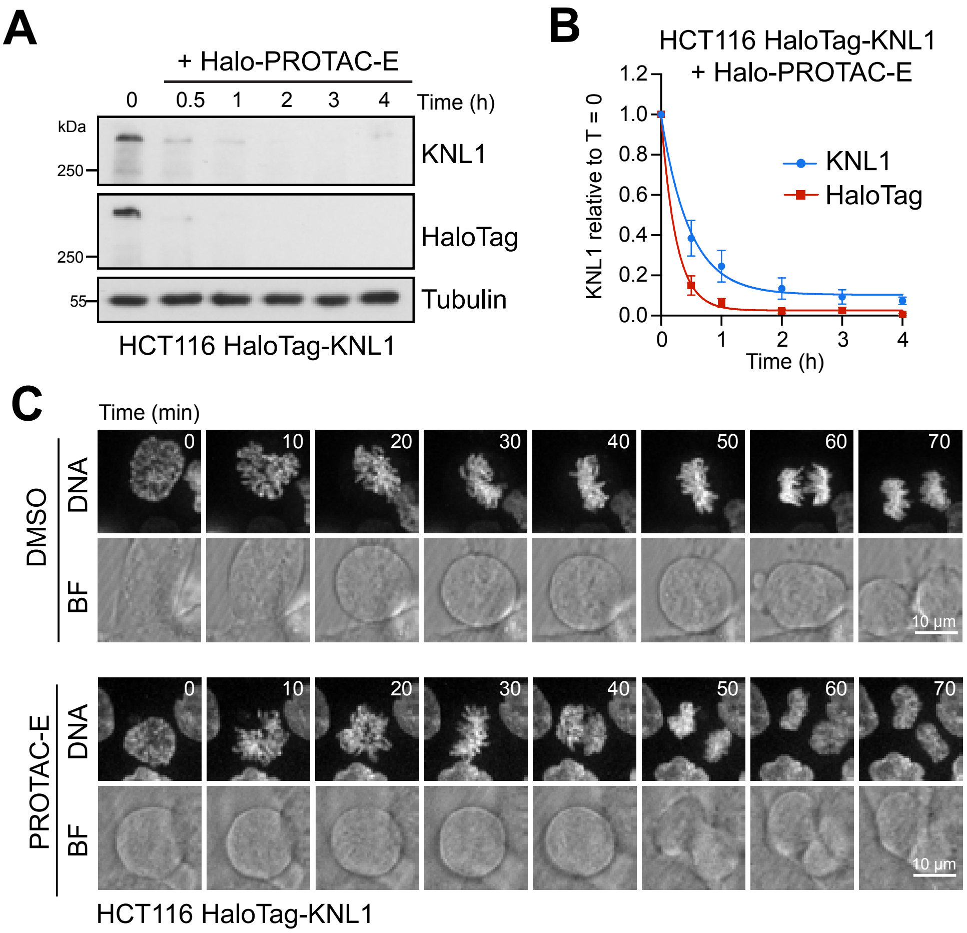
A
+ Halo-PROTAC-E
0 0.5 1 2 3 4 Time (h)
kDa
250
KNL1
250
HaloTag
55
Tubulin
HCT116 HaloTag-KNL1
B
HCT116 HaloTag-KNL1
+ Halo-PROTAC-E
KNL1 relative to T = 0
1.2
1.0
0.8
0.6
0.4
0.2
0.0
KNL1
HaloTag
0 1 2 3 4
Time (h)
C
Time (min)
DMSO
DNA
BF
0 10 20 30 40 50 60 70
10 μm
PROTAC-E
DNA
BF
0 10 20 30 40 50 60 70
10 μm
HCT116 HaloTag-KNL1

### Figure S4

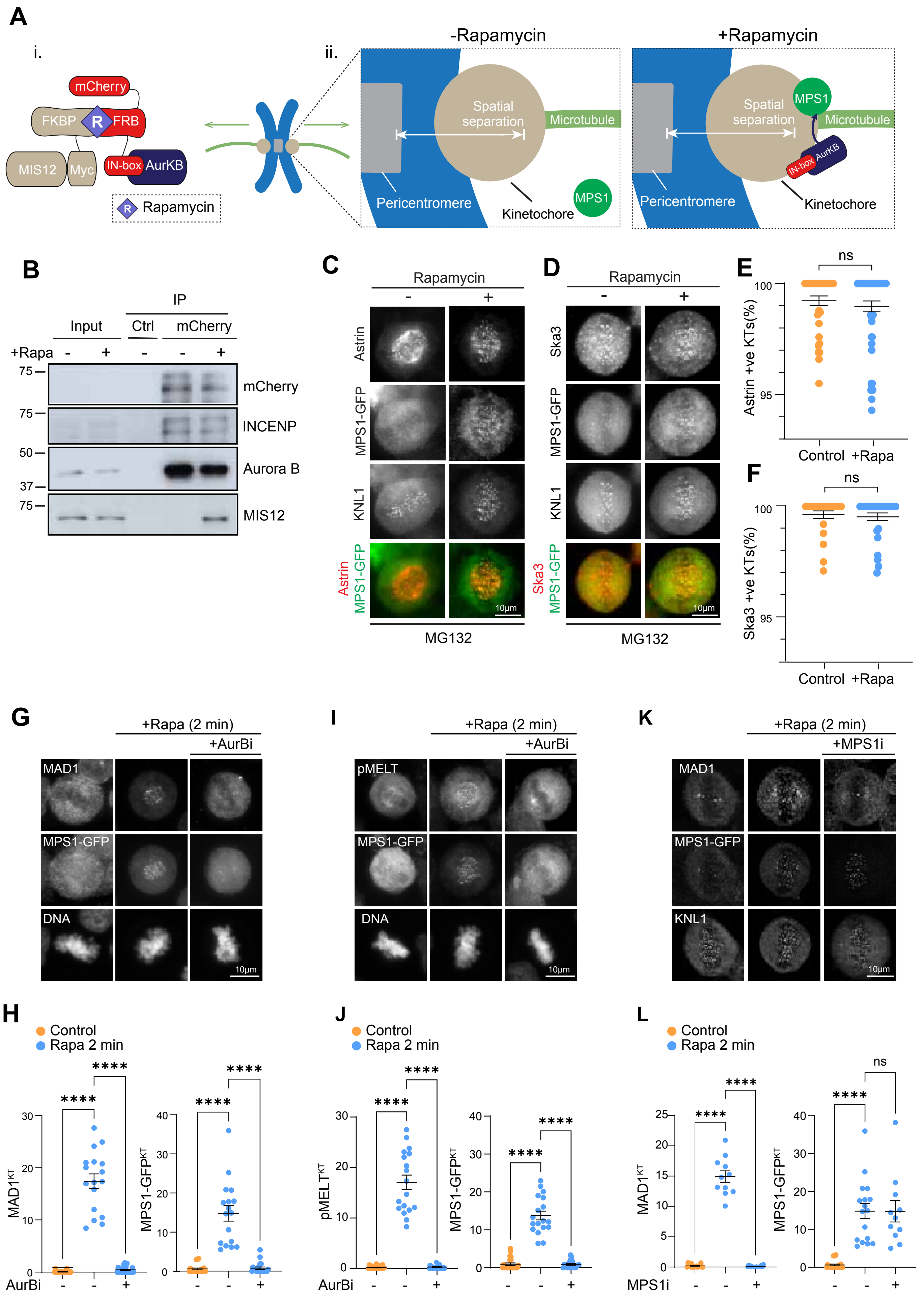

A
i.
mCherry
FKBP
R
FRB
MIS12
Myc
IN-box
AurKB
Rapamycin
ii.
-Rapamycin
Spatial separation
Microtubule
Pericentromere
Kinetochore
MPS1
+Rapamycin
B
IP
Input
Ctrl
mCherry
+Rapa
mCherry
INCENP
Aurora B
MIS12
C
Rapamycin
Astrin
MPS1-GFP
KNL1
10μm
MG132
D
Ska3
E
ns
Astrin +ve KTs(%)
Control
+Rapa
F
Ska3 +ve KTs(%)
G
+Rapa (2 min)
+AurBi
MAD1
DNA
I
pMELT
K
+MPS1i
H
Control
Rapa 2 min
AurBi
J
L
MPS1i
